# Biased gene retention in the face of massive nuclear introgression obscures species relationships

**DOI:** 10.1101/197087

**Authors:** Evan S. Forsythe, Andrew D. L. Nelson, Mark A. Beilstein

## Abstract

Phylogenomic analyses are recovering previously hidden histories of hybridization, revealing the genomic consequences of these events on the architecture of extant genomes. We exploit a suite of genomic resources to show that introgressive hybridization occurred between close relatives of Arabidopsis, impacting our understanding of species relationships in the group. The composition of introgressed and retained genes indicates that selection against incompatible cytonuclear and nuclear-nuclear interactions likely acted during introgression, while neutral processes also contributed to genome composition through the retention of ancient haplotype blocks. We also developed a divergence-based test to distinguish donor from recipient lineages without the requirement of additional taxon-sampling. Finally, to our great surprise, we find that cytonuclear discordance appears to have arisen via extensive nuclear, rather than cytoplasmic, introgression, meaning that most of the genome was displaced during introgression, while only a small proportion of native alleles were retained.

## Significance

Hybridization can lead to the transfer of genes across species boundaries, impacting the evolution of the recipient species through a process known as introgression (IG). IG can facilitate sharing of adaptive alleles but can also result in deleterious combinations of incompatible foreign alleles (i.e. epistatic incompatibility). How hybrids overcome these epistatic hurdles remains an open question. Here, we characterize IG in Arabidopsis and its closest relatives. Interestingly, our analyses favor an evolutionary scenario in which the vast majority of nuclear genes were displaced by foreign alleles during the evolution of *Capsella* and *Camelina*, obscuring species relationships. Simultaneously, a subset of nuclear genes resisted displacement, thereby minimizing epistatic incompatibilities between the organellar and nuclear genomes, suggesting one potentially fundamental mechanism for overcoming barriers to hybridization.

## Background

Hybridization is a driving force in plant evolution^1^, occurring naturally in ~10% of all plants, including 22 of the world’s 25 most important crops^2^. Botanists have long realized that through backcrossing to parents, hybrids can serve as bridges for the transfer of genes between species, a process known as introgression (IG). As more genome sequences become available, comparative analyses have revealed the watermarks of historical IG events in plant and animal genomes^3–5^. Cytonuclear discordance is a hallmark of many IG events, occurring, in part, because nuclear and cytoplasmic DNA differ in their mode of inheritance. In plants, this discord is often referred to as “chloroplast capture,” which has been observed in cases where IG of the chloroplast genome occurs in the near absence of nuclear IG or via nuclear IG to a maternal recipient6. Moreover, unlinked nuclear and cytoplasmic IG creates an interaction interface for independently evolving nuclear and cytoplasmic alleles, either of which may have accumulated mutations that result in incompatibilities with deleterious effects when they are united in hybrids. Such incompatibilities could exert a selective pressure that influences which hybrid genotypes are permissible thereby favoring the co-introgression of alleles for interacting genes7.

Disentangling IG from speciation is particularly important because IG may facilitate the transfer of adaptive traits. Robust statistical techniques^5, 8–15^ have been developed to detect the signatures of historical introgression (IG) in extant and extinct genomes. While existing techniques are able to identify the taxa that exchanged genes during IG using a four-taxon system, most methods do not explicitly distinguish which taxon served as donor and which as recipient during IG (i.e. polarization of IG directionality), an important distinction considering that IG impacts the evolution of the recipient lineage^4,6^. The existing methods that do polarize IG are only able to do so when there is a fifth taxon available, which diverged from its sister taxon involved in IG^11^, prior to the proposed IG event.

The wealth of genomic and functional data in Arabidopsis^16^, combined with publicly available genome sequence for 26 species make the plant family Brassicaceae an ideal group for comparative genomics. Phylogeny of the group has been the focus of numerous studies^17–23^, providing a robust estimate of its evolutionary history. While the genus *Arabidopsis* is well circumscribed^20,24^, the identity of its closest relatives remains an open question. Phylogenetic studies to date recover three monophyletic groups: clade A, including the sequenced genomes of *A. thaliana*^16^ and *A. lyrata*^25^; clade B, including the *B. stricta* genome^26^; and clade C, including the genomes of *Capsella rubella*, *C. grandiflora*^27^, and *Camelina sativa*^28^ (Supplementary Information). Analyses using nuclear markers strongly support A(BC), which is most often cited as the species tree^17,19,21–23^. Organellar markers strongly support B(AC)^18,19,29,30^ (Fig. 1a-b and Table S1). The genome sequences listed above can be used to explore the processes underlying this incongruence.

**Figure 1.**
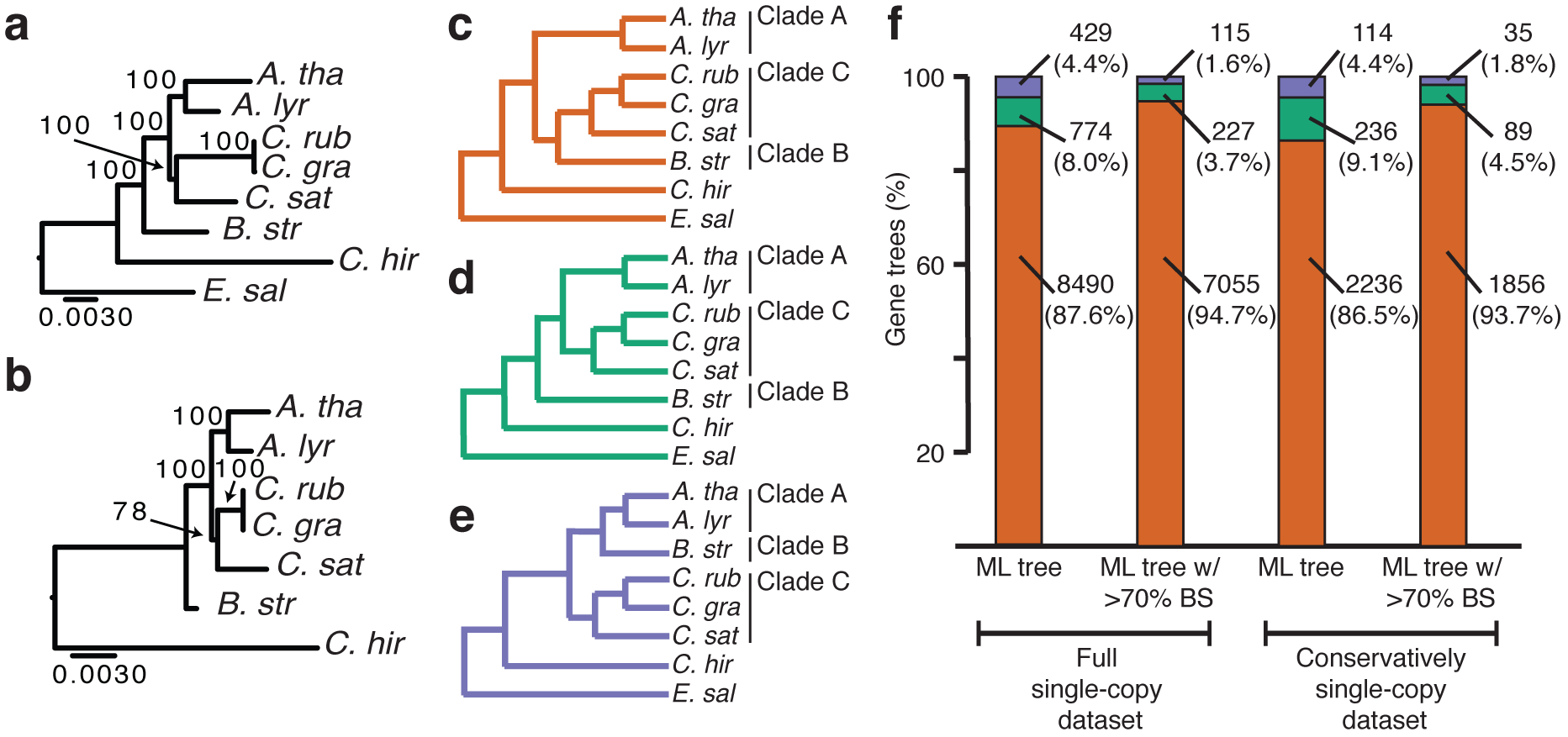
Incongruent gene tree topologies are observed within and between nuclear and organellar genomes. **a.** Chloroplast and **b.** mitochondria ML trees with branch support from 100 bootstrap replicates. Scale bars represent mean substitutions/site. **c-f.** ML gene tree topologies inferred from nuclear single-copy genes rooted by *E. salsugineum.* **c.** A(BC), **d.** B(AC) and **e.** C(AB) topologies. **f.** Numbers and frequencies of gene trees displaying A(BC) (orange), B(AC) (green), and C(AB) (purple). Single-copy genes are shown categorized by dataset and by level of bootstrap support.

Here, we exploit a suite of genomic resources to explore a putative chloroplast capture event involving Arabidopsis and its closest relatives by inferring gene trees for markers in all three cellular genomes from six available whole genome sequences. We document cytonuclear discordance and ask if it arose through IG of organelles or extensive IG of nuclear genes. Further, using a new divergence-based approach, we ask: Which lineage was the recipient of introgressed alleles? Finally, we explore the extent to which neutral processes, such as physical linkage as well as non-neutral processes, such as selection against incompatible alleles at interacting loci, shaped the recipient genome.

## Results

### Gene tree incongruence within and between organelle and nuclear genomes

We searched for incongruent histories present within and among nuclear and organellar genomes in representative species from each clade. We included *Cardamine hirsuta*^31^ and *Eutrema salsugineum*^32^ as outgroups. We considered three processes capable of producing incongruent histories: duplication and loss, incomplete lineage sorting (ILS), and IG. In addition, we assessed the possible contribution of phylogenetic error or ‘noise’.

Given the well-known history of whole genome duplication in Brassicaceae, we took extensive measures to minimize the possibility that duplication and loss biased our inferences. We identified single-copy nuclear genes as well as genes that were retained in all species post-duplication (see Discussion). In the chloroplast, we found 32 single-copy genes, while in mitochondria we identified eight. Maximum likelihood (ML) analyses of these yielded well-supported B(AC) trees (Fig. 1a and Fig. S2d-g). We identified 10,193 single-copy nuclear genes using *Orthofinder*^35^ (denoted as ‘full single-copy dataset’) (Fig. S1a-c). These genes were indicated as single-copy by *Orthofinder* because they form clusters that include exactly one locus from each species (with the exception of *C. sativa*, see **Methods**). These single-copy genes span the eight chromosomes of *C. rubella* (Fig. S1d), whose karyotype serves as an estimate of the ancestral karyotype for these species^36^. ML analyses yielded 8,490 (87.6%) A(BC), 774 (8.0%) B(AC), and 429 (4.4%) C(AB) trees (Fig. 1c-f and Table S2).

The most parsimonious explanation for our single-copy genes is that they were either not duplicated in our focal species or, if duplicated, were returned to single-copy before a speciation occurred, thus behaving as unduplicated in a phylogenetic context, meaning that any observed incongruent topologies resulted from a process other that duplication. However, while not parsimonious, it is important to consider the possibility that ancestral duplication, paralog retention through two speciation events, and lineage specific loss events led to hidden out-paralogs in our dataset. To further reduce the probability that this series of events contributed to incongruent gene trees, we further filtered our dataset to include only genes that were previously indicated as reliable single-copy markers in angiosperms^33,34^. This filter reduced our single-copy dataset to 2,098 genes (Fig. S1e-f). We combined this dataset with genes that were duplicated during whole genome duplication^37^ but did not undergo loss in focal species to yield a dataset of 2,747 genes, which we denote as ‘conservatively single-copy’, so named because they are the genes that are least likely to contain hidden out-paralogs. ML analyses of these genes yielded 2,236 (86.5%) A(BC), 236 (9.1%) B(AC), and 114 (4.4%) C(AB) trees (Fig. 1b-f), consistent with our results from the full single-copy dataset.

To ask whether phylogenetic noise contributed to incongruent nuclear gene tree topologies, we also filtered our single-copy nuclear gene tree results to contain only trees in which the observed topology was supported by at least 70% bootstrap support (BS) and found that B(AC) and C(AB) trees were still present (Fig. 1f). Together, these analyses confirm the incongruent histories present in the organellar and nuclear genomes and indicate that incongruence cannot be fully explained by gene duplication and loss or by phylogenetic noise.

### Contribution of introgression to incongruent gene trees

A number of approaches have been developed to determine the relative contributions of ILS and IG to gene tree incongruence. Comparative genomic approaches are based on the *D*-statistic^5,9^, which is typically applied to whole genome alignments and is calculated by determining the frequency of site patterns. It was not feasible to construct accurate whole genome alignments among our taxa, and thus we used multiple sequence alignments from single-copy genes to calculate *D-* and *F-*statistics. Analyses of both full and conservatively single-copy gene alignments indicated that introgression occurred (Table S3; positive *D* and *F*). Since phylogenomic analyses often focus on comparisons of gene trees rather than site-patterns, we also applied the rational of the *D-*statistic to gene trees, using gene tree topologies as proxies for site patterns to calculate a related statistic, referred to here as *D_GT_* (see **Methods**). Consistent with *D* and *F, D_GT_* indicated that ILS is sufficient to explain the frequency of C(AB) but not the observed frequencies of A(BC) and B(AC) in the nuclear genome (Table S4; positive *D_GT_*).

Coalescent based approaches^14,15^ use gene trees to distinguish between organismal histories that are tree-like (incongruencies among trees arise from ILS) and network-like (incongruencies result from ILS + IG). We analyzed our gene tree data in *PhyloNet*^15^ and found that reticulate networks were favored over tree-like evolution (Fig. S2j-q; ΔAIC ≥ 87.80 and ΔBIC ≥ 73.50). Similarly, *Tree Incongruence Checking in R* (*TICR*)^14^ indicated that a simple tree-like history fit the data poorly because the concordance factors for a significant proportion of quartets departed from expectation (Fig. S2r-u; *p* = 0.00058; *χ*^2^ test). In sum, both comparative genomic and coalescent based approaches support an evolutionary history that includes IG.

### Recovery of the species branching order and introgression events

To uncover which lineages were affected by IG, we determined the relative timing of the B(AC) and A(BC) branching events by calculating node depths (Fig. 2)^38^. IG nodes are expected to be younger than speciation nodes^12,38,39^ because IG produces incongruent trees when it occurs between non-sister species subsequent to speciation^4,5,9^ (illustrated by Fig. 2a). Therefore, we calculated the depth of the node uniting clade A with clade C in nuclear B(AC) trees and compared it with the depth of the node uniting the B and C clades in nuclear A(BC) trees (Fig. 2a-c, N.D.). We calculated node depths using four separate measures to account for potential biases (Fig. 2d-g). To account for selection on amino acids, we used synonymous divergence (*dS*) (Fig. 2d). To account for potential differing rates of evolution across the genome, we normalized *dS* using the divergence between the clade of interest and an outgroup (i.e. ‘relative node depth’)^12^ (Fig. 2e). To account for potential differences in rates of evolution between lineages, we also calculated node depths from ultrametric trees in which the rates of evolution had been smoothed across the tree using a penalized likelihood approach^40^ (Fig. 2f, Fig. S3, and Table. S5). To account for potential intragene discordance due to recombination within a gene, we divided each gene alignment into 200nt windows, inferred a neighbor joining tree for each window, and only calculated node depth from windows that were concordant with the ML tree for the gene, thus minimizing the probability of recombination within the loci from which node depth is calculated (Fig. 2g, Fig S4). For all four node depth measures, the node depth for A(BC) was significantly shallower than for B(AC) (Fig. 2d-g, Fig. S3, Fig S4 and Table S6; *p*<2.2e-16, Wilcoxon), indicating that IG rather than speciation produced the observed A(BC) nuclear gene trees. This result is insensitive to the removal of the deepest nodes in both A(BC) and B(AC) bins (Fig. S3o-t). Hence, node depth data suggest that A and C diverged from each other prior to the exchange of genes between clade B and C via IG. This surprising result stands in opposition to previously published trees inferred from single or concatenated nuclear genes, which strongly favor A(BC)^20, 22–24^. However, it bolsters the argument that B(AC) best represents the species branching order despite the low frequency of these genes in the nucleus (similar to ^38^), and further suggests that the vast majority of nuclear genes in either B or C arrived there via IG. We discuss the implications of this finding on the concept of the species branching order (see **Discussion**). It should be noted that our downstream analyses of selection and neutral processes (Fig. 4, Fig. S6, and Table S6) are framed in the context of nuclear introgression but would remain equally valid if cytonuclear discordance arose via organellar introgression.

**Figure 2.**
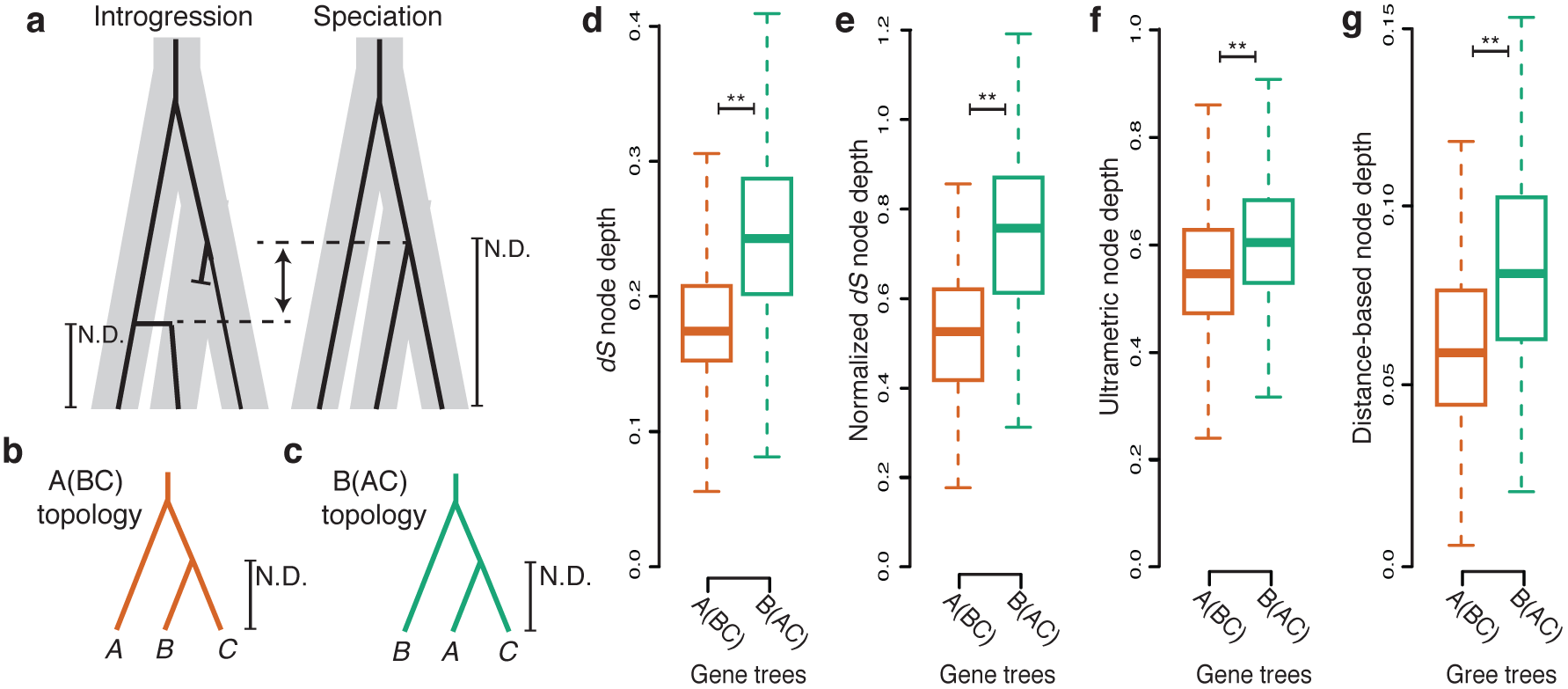
Node depths indicate extensive introgression led to transfer of nuclear genes. **a.** Model depicting expected node depths (N.D.) for genes undergoing IG (left) or speciation (right). Speciation history is represented by thick grey bars. Individual gene histories are represented by black branches. Blunt ended branches represent a native allele that was replaced by an IG allele. Vertical arrow indicates expected difference in node depth. **b-c.** The informative node depths on A(BC) (**b**) and B(AC) (**c**) trees. **d-f.** Boxplots depicting observed median and quartile node depths measured from *dS* (**d**), normalized *dS* (**e**), ultrametric gene trees (**f**), and concordant windows within gene alignments (**g**).

### Identification of unidirectional introgression donor and recipient linages

We next asked whether transfer of genetic material during IG was unidirectional and, if so, which of the two clade ancestors was the donor and which was the recipient of introgressed alleles. Existing methods for polarizing the direction of IG require additional taxa with specific phylogenetic positioning relative to the introgression event^5,11^. No such taxa are available for our inferred introgression event; therefore, existing polarization methods are not applicable to our data. Instead, we present a divergence-based approach to infer directionality of IG, calculated from pairwise sequence divergence between taxa involved in IG and a sister taxon by comparing divergence values obtained from introgressed loci *vs*. non-introgressed loci (see **Methods**).

We calculated the rate of pairwise *dS* for all pairs of species and used these to determine the average *dS* between pairs of clades (B vs. C = *dS(B,C)*; A vs. C = *dS(A,C)*; A vs. B = *dS(A,B)*) (Fig. S5). We denoted *dS* values with *_SP_* when obtained from B(AC) trees (our inferred species branching order) and *IG* when obtained from A(BC) trees (IG branching order) (Fig. 3a and b). We compared *dS(B,C)_IG_*, *dS(A,C)_IG_*, and *dS(A,B)_IG_* to *dS(B,C)_SP_*, *dS(A,C)_SP_*, and *dS(A,B)_SP_*, respectively, to ask if divergence is consistent with unidirectional IG from B to C (Fig. 3a) or from C to B (Fig. 3b), or with bidirectional IG. We found that *dS(B,C)_SP_* > *dS(B,C)_IG_* (*p*<2.2e-16, Wilcoxon), *dS(A,C)_SP_* < *dS(A,C)_IG_* (*p*=2.365e-12), and *dS(A,B)_SP_* = *dS(A,B)_IG_* (*p*=0.1056), indicating unidirectional IG from clade B to clade C (Fig. 3c and Fig. S5). This result is consistent with the *Phylonet* network shown in Fig. S2m and one shown in Fig. S2n, which respectively indicate that 96.6% and 90.5% of sampled nuclear alleles were introgressed from clade B to C.

**Figure 3.**
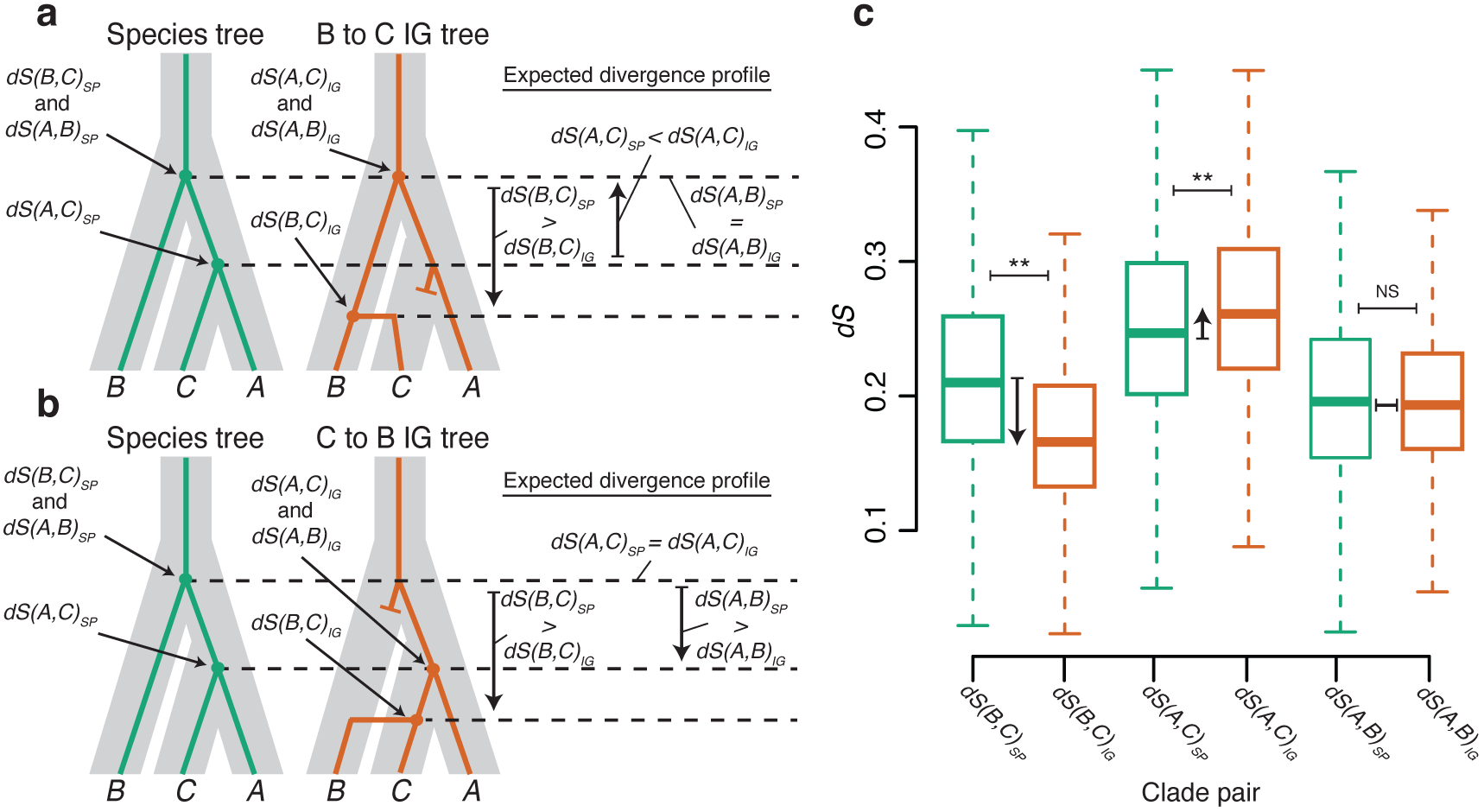
Unidirectional introgression led to transfer of nuclear genes from clade B to clade C. **a-b.** Model depicting pairwise *dS* divergence between clades A, B, and C. Arrows point to nodes on the species tree (B(AC)) and the IG tree (A(BC)) indicated with *SP* and *IG* subscripts, respectively. Expected node depths under IG from clade B to clade C **(a)** or from clade C to B **(b)**. Vertical arrows depict expected differences between gene trees representing speciation and IG. **c.** Observed *dS* distances on speciation gene trees (green boxes; *dS(B,C)_SP_, dS(A,C)_SP_,* and *dS(A,B)_SP_*) and IG gene trees (orange boxes; *dS(B,C)_IG_, dS(A,C)_IG_,* and *dS(A,B)_IG_*). Arrows indicate observed differences between *SP* and *IG* comparing *dS(B,C), dS(A,C),* and *dS(A,B)*. Horizontal bars above boxes in **c** represent distribution comparisons. \*\**p*<0.01, *NS p*>0.05.

### The role of cytonuclear interactions during introgression

The IG that occurred during the evolution of clade C resulted in a genome in which the majority of nuclear alleles were displaced by alleles from clade B, while native organellar genomes were maintained. We asked whether we could detect patterns within the set of nuclear genes that were also maintained alongside organelles during IG. We hypothesized that during the period of exchange, selection would favor the retention of alleles that maintain cytonuclear interactions, especially when replacement with the paternal allele is deleterious^7^. Using Arabidopsis Gene Ontology (GO) data^41^, we asked if B(AC) nuclear genes were significantly enriched for chloroplast and mitochondrial-localized GO terms, indicating that these genes are more likely to be retained than are other nuclear genes. We calculated enrichment (*E*) for each GO category by comparing the percentage of B(AC) nuclear genes with a given GO term to the percentage of A(BC) genes with that term (see **Methods**). Positive *E* indicates enrichment among B(AC) genes; negative *E* indicates enrichment among A(BC) genes. B(AC) nuclear genes are significantly enriched for chloroplast (*E*=0.10, *p*=0.00443, 1-tail Fisher’s) and mitochondrial localized (*E*=0.13, *p*=00250) GO terms (Fig. 4a and Table S6). Enrichment was also detected at the level of organelle-localized processes such as photosynthesis (*E*=0.29, *p*=0.01184), including the light (*E*=0.44, *p*=0.00533) and dark (*E*=0.65, *p*=0.04469) reactions. The opposite enrichment pattern exists for nuclear localized genes (*E*=-0.06, *p*=0.00936) (Fig. 4a). In sum, these results suggest a role for selection in shaping which genes were displaced during IG.

**Figure 4.**
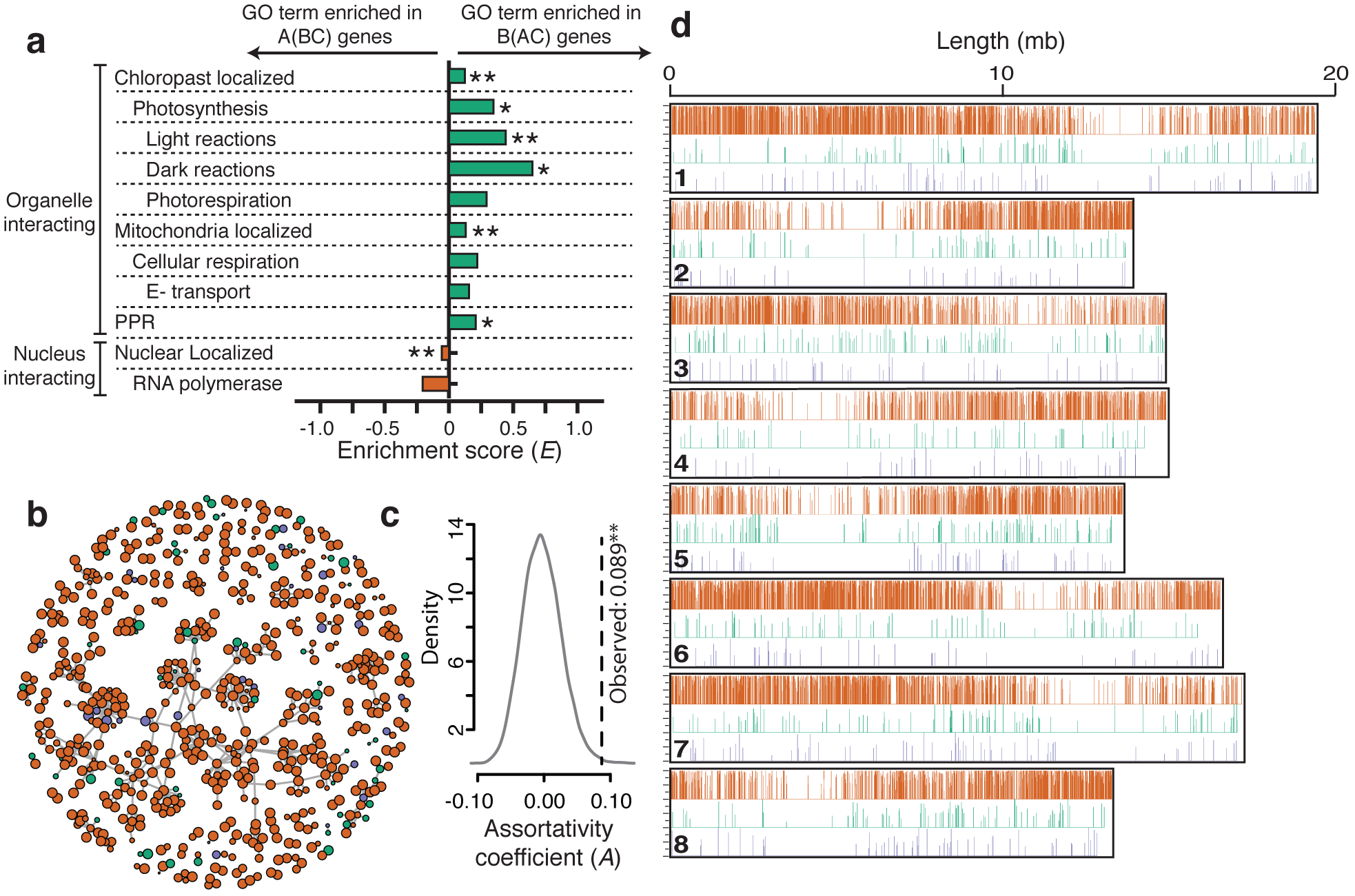
The genomic consequences of epistasis and genetic linkage during IG. **a.** Enrichment (*E*) for GO terms = (% B(AC) genes – % A(BC) genes) / (% B(AC) + A(BC) genes). **b.** Protein-protein interaction network for Arabidopsis protein complexes. Node fill, gene tree topology; node diameters proportional to bootstrap support (Fig. S2a-c). **c.** Assortativity coefficient (*A*) of the network. Null distribution of *A* (grey curve); dotted line, observed *A.* **d.** Nuclear genes mapped to *C. rubella*. Vertical lines, genes (colored by topology). Line heights proportional to bootstrap support (Fig. S2a-c). \*\**p*<0.01, \**p*<0.05.

### The role of nuclear-nuclear interactions during introgression

We also asked if interactions between/among nuclear genes influenced the likelihood of replacement by foreign alleles. Using Arabidopsis protein-protein interaction data^42^, we constructed an interaction network of the full set of single-copy nuclear genes (Fig 4b). To assess whether genes with shared history are clustered in the network, we calculated its assortativity coefficient (*A*) (**Methods**). We assessed significance by generating a null distribution for *A* using 10,000 networks of the same size and shape with randomized topology assignments. In our empirical network, *A* was significantly positive (*A*=0.0885, *p*=0.00189, *Z-*test), and hence topologies are clustered (Fig. 4c), indicating that selection acted against genotypes containing interactions between maternal and paternal alleles.

### The role of physical linkage during introgression

While it appears gene function exerted influence on nuclear IG, we also wondered whether blocks of genes with similar histories were physically clustered on chromosomes. We looked for evidence of haplotype blocks using the *C. rubella* genome map (Fig. 4d). Previous studies in this group estimate linkage disequilibrium to decay within 10kb^43,44^, creating blocks of paternal or maternal genes around that size. We assessed the physical clustering of genes with shared history by two measures: 1) number of instances in which genes with the same topology are located within 10kb of each other (Fig. S6a), and 2) number of instances in which neighboring genes share topology, regardless of distance (Fig. S6b). The second measure provides a simple measure of clustering without requiring an estimate of ancestral linkage. We compared both measures to a null distribution generated from 10,000 replicated chromosome maps in which the topology assignments were randomized across the marker genes. By both measures, we found significant clustering of A(BC) (measure 1: *p*=3.022e-8; measure 2: *p*=1.41364e-10, *Z*-test) and B(AC) (measure 1: *p*=0.003645; measure 2: *p*=1.7169e-11) genes (Fig. S6c-h). The observed clustering indicates that haplotype blocks of co-transferred and un-transferred genes are detectable in extant genomes, pointing to physical linkage as a factor influencing whether genes are transferred or retained.

## Discussion

Phylogenomic studies in plants face unique challenges. The prevalence of gene and genome duplication complicates the detection of orthologs, and thus choosing markers that minimize duplication is extremely important when applying tests of IG originally developed for animals5. Since duplication history cannot be definitively known, we can never be sure that cryptic duplication has not introduced phylogenetic incongruence into our dataset; this is a risk in any phylogenetic study, especially in plants. We acknowledge that all nuclear genes have undergone duplication at some point in Brassicaceae^37^ and address this challenge by specifically targeting genes least likely to have undergone duplication during the speciation and introgression events we detected. If duplication was biasing the results we obtained from our full single-copy dataset, we expected that the proportion of B(AC) trees would have decreased in our conservatively single-copy dataset. However, the proportions we observed were not substantially impacted by our conservative single-copy filter. In fact, the proportion of B(AC) genes was slightly higher in the conservatively single-copy genes, the opposite of what we would expect if duplication was creating incongruent trees. Moreover, results of the *D*-, *F-,* and *D_GT_-*statistics from both datasets significantly indicated IG (Table S3, and Table S4), another indication that biases associated with cryptic duplication and loss are not driving our conclusions of IG.

We applied several methods to distinguish between IG and ILS. Like all applications of *D* and related statistics, it’s important to acknowledge that ancestral population structure may produce signatures that mimic IG^45^. However, when this possibility was thoroughly explored in the case of Neanderthal, IG remained the favored hypothesis^46^. Here, regardless of the measure or approach employed, our results (Fig. S2, Table S3, and Table S4), were always consistent with an explanation of IG rather than ILS or duplication and loss. While we appreciate the limitations of each approach, here we argue that the consistent finding of IG favors this hypothesis over all others.

Our initial interpretation of the observed phylogenetic incongruence was that A(BC) resulted from simple speciation events and B(AC) resulted from IG between clades A and C, a pattern we referred to as cytoplasmic IG. However, in light of recent findings from mosquitos^38,47^, we thought it important to consider alternative hypotheses. Using the same approach that revealed IG in mosquitos, we calculated the mean node depth for each of the alternative topologies we recovered for nuclear genes. In addition, we employed several strategies to account for the effects of selection (Fig. 2d), effective population size variation across the genome (Fig. 2e), linage-specific effects (Fig. 2f). and intragenic recombination (Fig. 2g) on our node depth calculations. In all cases, our node depth comparisons rejected the hypothesis that the node uniting clades A and C on B(AC) trees resulted from an introgression event, and instead indicated that the node uniting clades B and C on A(BC) trees resulted from an introgression between clades B and C. Based on these results, we suggest that the ‘true’ species branching order is B(AC).

There is growing debate about the efficacy of bifurcating phylogenies in describing organismal evolution, prompting the development of powerful network frameworks that highlight reticulation in species relationships. While our analysis reinforces the importance of considering reticulation, we also show that bifurcating trees should not be entirely abandoned in the face of reticulation. The presence of reticulation does not preclude the occurrence of simple bifurcating speciation events, it simply means some bifurcations result from speciation while others result from IG. Therefore, some gene trees will have nodes representing speciation events while other genes trees will have a node or nodes that represent IG. We define the ‘true’ species branching order as the topology of the gene tree in which all nodes represent speciation events, even if this history does not represent the majority of the genome. Our finding of massive nuclear IG leads to a dilemma regarding which branching order should be used in future comparative studies in this group. For many (if not most) practical purposes, it is reasonable to continue to use A(BC) because it represents the history of most of the genome. However, studies using this topology should bear in mind that this history is more complicated than simple speciation and consider the potential implications. Integrating all available information into a useful model for studying trait evolution represents a future goal in systematics.

We demonstrate the use of several complementary techniques to identify the taxa that exchanged genes during IG, many of which operate in a four-taxon (or four-clade) context. However, most methods do not explicitly distinguish which taxon served as donor and which as recipient during IG. The existing methods that do polarize IG are only able to do so when there is a fifth taxon (or clade)^11^. The divergence-based approach presented here can be applied to infer the directionality of IG in a four-taxon case when additional taxa are not available. It should be noted that our goal in the present study was to present the conceptual framework of divergence-based polarization of IG and to lay the groundwork for further development of these types of methods. It was not our goal, here, to mathematically derive the test or to explore parameter robustness. For example, factors such as population size and structure, divergence time, size of loci, rate of evolution^48^, and extent of linkage disequilibrium^49^ have been demonstrated to affect existing statistics for inferring IG^9,45^ but have not been explored here. We have also not explored the power of our test to polarize IG when it is asymmetrical but not strictly unidirectional, all of the above representing important next steps toward understanding the conditions under which divergence-based phylogenetic methods can accurately recover the direction of IG.

Applied to genomic data, our test infers IG of nuclear genes from clade B to clade C. Since cytoplasmic inheritance is matrilineal in Brassicaceae, we conclude that clade C was the maternal recipient of paternal clade B nuclear alleles. While we can only postulate about the specific crosses and backcrosses that occurred during IG, it is likely that F1 hybrids arose from a clade C maternal parent and clade B paternal parent. We find evidence that selection acted during the backcrosses that followed, resulting in resistance of organelle interacting nuclear genes to replacement by paternal alleles. Maternal nuclear alleles that function in chloroplasts or mitochondria in fundamental processes were not replaced at the same rate as maternal alleles localized to other areas of the cell or for other functions. These genes may constitute a core set whose replacement by paternal alleles is deleterious. We also find evidence that selection acted to maintain nuclear-nuclear interactions. In general, our results suggest that epistatic interactions between genes exerted selective pressure that influenced which genes were displaced and which were retained. Whether this type of selection drove the displacement or retention of entire haplotype blocks via hitchhiking remains a future question.

In summary, our comparative genomic analyses are consistent with an evolutionary history in which massive unidirectional nuclear IG, driven by selection and influenced by linkage, underlie the original observation of “chloroplast capture.” The species branching order in this group is more accurately reflected by B(AC), and thus similar to the findings of ^38^, nuclear IG obscured speciation such that the latter was only recoverable from extensive genomic data. What makes IG here particularly interesting is that its impact on the genome is evident despite the fact that it must have occurred prior to the radiation of clade A 13 – 9 million years ago^20,22^. Hence, it’s likely that, as additional high-quality genomes become available, comparative analyses will reveal histories that include nuclear IG, even when the genomes considered are more distantly related. In short, our findings explore the genomic battle underlying chloroplast capture to reveal an onslaught of alleles via unidirectional IG. A core set of nuclear genes resisted displacement by exogenous alleles; purifying selection removed genotypes with chimeric epistatic combinations that were deleterious, just as Bateson-Dobzhansky-Muller first described^7,50^. Will other IG events reveal similar selective constraints as those we detail? If so, it could point us toward key interactions between cytoplasmic and nuclear genomes that lead to successful IG, thereby refining our understanding of the factors governing the movement of genes among species.

## Methods

### Phylogenomic pipeline

#### Clustering of putative orthologs

Coding sequences (CDS) for *Arabidopsis thaliana*, *A. lyrata*, *Capsella rubella*, *C. grandiflora*, *Boechera stricta*, and *Eutrema salsugineum* were obtained from *Phytozome*16,25–27,32,51; *Camelina sativa* and *Cardamine hirsuta* were obtained from *NCBI* 28,31. Datasets were processed to contain only the longest gene model when multiple isoforms were annotated per locus. CDS were translated into amino acid (AA) sequences using the standard codon table. The resulting whole proteome AA sequences for the eight species were used as input to cluster orthologs via *Orthofinder* (version 1.1.4)^35^ under default parameters (Fig. S1a). Two different filtering strategies with varying stringency were applied to the resulting clusters to yield two dataset partitions referred to as ‘full single-copy dataset’ and ‘conservatively single-copy dataset’. Both filtering strategies are described below.

#### Full single-copy dataset filtering

The full single-copy dataset was identified by sorting *Orthofinder* results to include only clusters that contained exactly one sequence per species, except in the case of *C. sativa*. Clusters containing one to three sequences from *C. sativa* were also retained as single-copy (Fig. S1b) because it is a hexaploid of relatively recent origin. Thus, clusters with up to three *C. sativa* paralogs (*i.e*. homeologs) were retained, and we expected these homeologs to form a clade under phylogenetic analysis (see **Multiple sequence alignment and gene tree inference of nuclear genes**). Gene clusters that yielded trees deviating from this expectation were omitted from further analysis. The full single-copy dataset also contains groups classified as retained duplicates (Fig. S1c). Retained duplicate clusters contain exactly two sequences per species (three to six in *C. sativa*). The *A. thaliana* sequences in each cluster represent known homeologs from the *α* whole genome duplication that occurred at the base of Brassicaceae^37^, and thus is shared by all sampled species in this study. We retained only those gene clusters that produced trees in which the paralogs formed reciprocally monophyletic clades (Fig. S1c).

#### Conservative single-copy dataset filtering

We also used a more stringent set of criteria to develop a conservatively single-copy dataset. For this dataset, we compared the results obtained from *Orthofinder* with results from previously published assessments of plant single-copy or low copy gene families^33,34^. The criteria and taxon sampling of our *Orthofinder* filtering and the filtering strategies of the two previous analyses differed, meaning each analysis provides its own level of stringency. Moreover, both previous analyses included *A. thaliana*, allowing for direct comparison with our results. We filtered our clusters to include only those genes recovered by both *Orthofinder* and in at least one published analysis. We refer to these as conservatively single-copy. Conservatively single-copy genes plus the retained duplicates described above constitute the conservatively single-copy dataset. CP and MT gene datasets were filtered using the same criteria used to filter the full single-copy dataset.

#### Multiple sequence alignment and gene tree inference of nuclear genes

For single-copy genes, we generated AA-guided multiple sequence alignment of CDS using the *MAFFT* algorithm (version 6.850)^52^, implemented using *ParaAT* (version 1.0)^53^, under the default settings for both. Multiple sequence alignments of CDS for each gene cluster were used to infer maximum likelihood gene trees using *RAxML* (version 8)^54^ under the general time reversible model with gamma distributed rate heterogeneity. Support values for nodes were calculated from 100 bootstrap replicates using rapid bootstrapping.

#### Assembly and annotation of mitochondria and chloroplast genomes

Whole genome sequence reads for *A. lyrata*, *B. stricta*, *C. rubella*, *C. grandiflora,* and *C. sativa* were acquired from *NCBI’s Sequence Read Archive* (SRA). The run IDs of SRA files used to assemble organelle genomes for each species were: *A. lyrata* (DRR013373, DRR013372); *B. stricta* (SRR3926938, SRR3926939); *C. rubella* (SRR065739, SRR065740); *C. grandiflora* (ERR1769954, ERR1769955); *C. sativa* (SRR1171872, SRR1171873). Both SRAs for each species were independently aligned to the *Arabidopsis thaliana* mitochondrial (MT) genome (Ensembl 19) using *HiSat2*^55^ with default settings for paired-end reads within *CyVerse’s Discovery Environment*^56^. 15-30X coverage was recovered for each alignment. Mapped read alignment files were converted from BAM to SAM using *SAMtools*^57^. MT consensus sequences were generated (base pair call agreement with 75% of all reads) from each alignment within *Geneious* (version 7.0; Biomatters)^58^. Each MT consensus sequence was annotated based on the *A. thaliana* MT genome annotation (Ensembl 19). CDSs were then extracted using gffread from the *Cufflinks* package^59^. The same method was used to assemble the *B. stricta* CP genome. All other chloroplast genome sequences were publicly available.

#### Multiple sequence alignment and tree inference from chloroplast and mitochondria markers

Single-copy CP and MT genes were identified, aligned, and used to infer phylogeny as described previously for nuclear genes. Individual gene tree results are presented in Fig. S2d-e. We also generated concatenated alignments for both the CP and MT genes using *SequenceMatrix*^60^. We inferred trees (Fig. 1a-b) from both concatenated alignments using *RAxML* with the same parameters described above.

### Downstream analyses

#### Gene tree topology analysis

Tree sorting was performed in batch using the *R* packages, *Ape*^61^, *Phangorn*^62^, and *Phytools*^63^. Gene trees from the retained duplicates were midpoint rooted and split at the root into two subtrees, each of which contained a sequence from all eight analyzed species. Subtrees were analyzed as individual trees alongside all other single-copy gene families as described below. First, each gene tree was rooted at *E. salsugineum*. Next trees were sorted by considering the topological arrangement of the A, B, and C lineages. For example, a tree was categorized A(BC) if *B. stricta*, *C. rubella*, *C. grandiflora*, and *C. sativa* formed a monophyletic clade. Thus, the branch in the tree leading to the monophyletic clade (the branch uniting *B. stricta*, *C. rubella*, *C. grandiflora*, and *C. sativa* in the above example) was considered the topology-defining branch. Statistical support for any given tree was summarized as the bootstrap value along the topology-defining branch.

Since the focus of our analysis was on topological incongruence of A, B, and C clades, our topology assessment was not designed to detect topological arrangements within A, B, and C clades or in other parts of the trees. If a gene cluster failed to form either a monophyletic A or C clade following phylogenetic analysis, it was marked as ‘other topology’ and removed from further downstream analysis. Exact topologies of all trees, including those recorded as ‘other topology’, are summarized in Table S2.

#### Applying *D*, *F*, and *D_GT_* statistics to assess the effects of incomplete lineage sorting and introgression

To determine whether the observed gene tree incongruences could have been caused primarily by incomplete lineage sorting (ILS), we calculated Patterson’s *D*-statistic (*D*) (also known as the ABBA-BABA or 4-taxon test)^5,9^. *D* is typically applied to whole genome alignments of three in-group taxa and one out-group taxon. It is calculated by scanning the alignment to identify site patterns consistent with two possible resolutions of ILS (ABBA and BABA). Due to the relatively deep divergence and numerous chromosomal rearrangements between genomes used here, it was not feasible to construct accurate whole genome alignments. Instead, we identified ABBA and BABA site patterns within single-gene multiple sequence alignments used to infer gene trees. We calculated *D* and *F* using the total number ABBA and BABA sites from all nuclear gene alignments (or subsets of nuclear genes corresponding to individual chromosomes or conservatively single-copy genes). We excluded *C. sativa* sequences from this analysis due to the presence of multiple *C. sativa* paralogs in some trees. We considered only biallelic sites in which the two outgroups, *E. salsugineum* and *C. hirsuta,* have the same allele. We also required individual species within each clade to have the same allele. For example, an ABBA site would be one in which *E. salsugineum, C. hirsuta, A. thaliana, A. lyrata, C. rubella, C. grandiflora,* and *B. stricta* display T, T, G, G, G, G, and T, respectively. Note that all members of clade A and C share the derived allele. An example of a BABA site would be T, T, G, G, T, T, and G, respectively. In this case, members of clades A and B share the derived allele. We also tallied AABB sites, (e.g. T, T, T, T, G, G, and G, respectively), in which clades B and C share the derived allele, although AABB sites are not a component of *D* or *F.* We calculated *D* and *F* according to the equations from^64^. All site counts and statistics are shown in Table S3.

We also applied the rationale of *D* to gene tree topology counts by calculated a related statistic, *D_GT_.* We used gene tree topologies as proxies for site patterns. Since B(AC) and C(AB) trees were closest in frequency in the nuclear genome, we asked whether their frequencies were statistically significantly different using *D_GT_*. B(AC) trees and C(AB) trees were treated as ABBA and BABA sites, respectively, while A(BC) was treated as AABB. *D_GT_* was then calculated as follows:

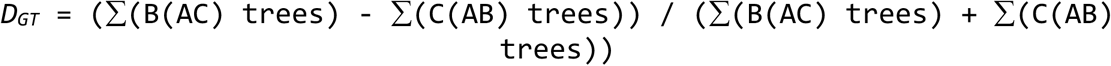

We calculated *D_GT_* for the set of all nuclear genes as well as for subsets of genes present on each of *C. rubella’s* nuclear chromosomes^36^. Results from all *D_GT_* calculations are given in Table S4.

#### Phylogenetic network reconstruction and introgression analysis

To evaluate the likelihood that the observed incongruence was caused by IG, we also reconstructed maximum likelihood phylogenetic networks using InferNetwork_ML in *PhyloNet* (version 3.6.1)^15^. We input all nuclear gene trees (Fig. S1d, *Full single-copy genes* dataset) and implemented InferNetwork_ML using the command ‘InferNetwork_ML (all) *h* –n 100 –di –o –pl 8;’, where *h* is the number of reticulations allowed in a given network. The method ignores gene tree branch lengths, utilizing gene tree topologies alone to infer reticulation events. We performed separate analyses using *h* = 0 (a tree), *h* =1, and *h* = 2, outputting the 100 most likely trees/networks (designated with –n) from each analysis. We followed the analysis strategies of^65^, manually inspecting networks to identify those with edges consistent with both the major nuclear topology [A(B,C)] as well as the major CP and MT topology [B(A,C)] (Fig. S2l-o). Additionally, we reported the most likely tree/network from each analysis (Fig. S2k, p-q). As an additional means of asking whether ILS alone adequately explains incongruence, we performed Tree Incongruence Checking in R (TICR)14. We used a population tree inferred from *PhyloNet* (*h* = 0) (Fig. S2j) with a table of concordance factors for all quartets. We performed the *TICR* test as implemented in the *R* package, *phylolm*^66^, according to the methods outlined in: https://github.com/crsl4/PhyloNetworks.jl/wiki/TICR-test:-tree-versus-network%3F.

#### Identification of introgressed topology and species branching order

In order to identify the topology most likely to represent IG, we measured node depths on trees displaying the A(BC) B(AC). As above, *C. sativa* sequences were not considered in order to avoid complications associated with paralogous sequences. For each nuclear gene tree, we calculated pairwise synonymous divergence (*dS*) between taxa on the tree using *PAML* (version 4.8)^67^. To infer the pairwise distance between two clades on the tree, we took the average *dS* score between each combination of taxa present in the two clades. For example, the depth of the node uniting clades A and C on B(AC) trees would be the average of *dS*(*A. thaliana*, *C. rubella*), *dS*(*A. lyrata*, *C. rubella*), *dS*(*A. thaliana*, *C. grandiflora*), and *dS*(*A. lyrata*, *C. grandiflora*). To calculate normalized *dS,* each *dS* node depth (as described above) was divided by the average pairwise *dS* of each ingroup species versus the outgroup, *C. hirsuta.*

We also calculated node depths from ultrametric gene trees. Before measuring node depths, gene trees were smoothed to ultrametric trees using semiparametric penalized likelihood rate smoothing^40^. We implemented the rate smoothing algorithm designated by the *chronopl* function in the *Ape* package. We tested six values of the smoothing parameter (λ), which controls the tradeoff between parametric and non-parametric formulation of rate smoothing, to assess the sensitivity of node depths to different values of λ. We calculated node-depth on ultrametric trees for nodes representing *T_1_* and *T_2_* on each given topology (Fig. S3a). We plotted the frequency distributions of node depths (Fig, S3b) as well as descriptive statistics (Fig. S3c-t).

In order to account for intragenic recombination, we split each gene alignment into 200nt alignments, the goal being to reduce the probability of recombination occurring in the middle of our alignment. For each window, we calculated a distance matrix and inferred a neighbor joining “window tree” using *Ape* in *R*^61^. We calculated the depth of the *T_1_* node for each window displaying either A(BC) or B(AC) from the distance matrix by averaging the pairwise distance values similar to our treatment of *dS* node depths above. We documented the number of discordant windows in alignments for A(BC) (Fig. S4a) and B(AC) (Fig. S4b) trees and used boxplots to compare distributions of A(BC) and B(AC) node depths (Fig. 2g and Fig. S4c).

#### Divergence-based polarization of introgression

For each nuclear gene tree from our Brassicaceae dataset, we calculated pairwise synonymous divergence (*dS*) between taxa on the tree using *PAML* (version 4.8)^67^. To infer the pairwise distance between two clades on the tree, we took the average *dS* score between each combination of taxa present in the two clades. We excluded *C. sativa* sequences from this analysis due to the presence of multiple *C. sativa* paralogs in some trees. We define *dS* between clades B and C, clades A and C, and clades A and B as *dS(B,C)*, *dS(A,C),* and *dS(A,B)*, respectively (Fig. 3 and Fig. S5). For example, to calculate the distance between clade A and clade C (*dS(A,C)*) for a given tree, we used the following equation:

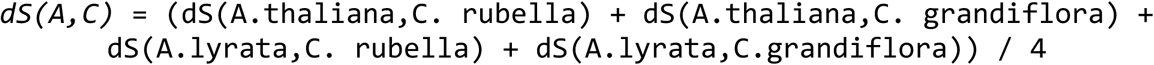

We calculated *dS(X,Y)* for both the species branching order, B(AC), and the introgression tree, A(BC) (*dS(X,Y)_SP_* vs. *dS(X,Y)_IG_,* respectively). Frequency distributions of each value were determined.

#### GO category enrichment analysis

*Gene Ontology* (GO)^41^ data for Arabidopsis were obtained from *The Arabidopsis Information Resource* (www.arabidopsis.org)^16^. We determined the GO terms associated with the Arabidopsis genes present in our full single-copy data set. For each GO term, the percentage of B(AC) trees containing the GO term was compared to the percentage of A(BC) trees containing it. Comparisons were quantified with an enrichment score (*E*). For example, we used the following equation to ask if B(AC) or A(BC) topology genes are enriched for CP localization:

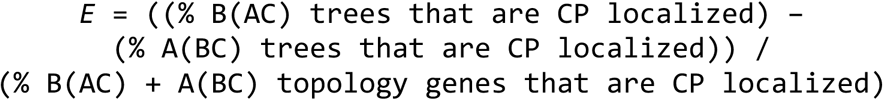

Positive *E* indicates enrichment for a given GO category among B(AC) trees, while negative *E* indicates enrichment among A(BC) trees (Table S6).

#### Network analysis of protein-protein interactions

Experimentally curated protein-protein interaction data for Arabidopsis were downloaded from *Arabidopsis thaliana Protein Interaction Network (AtPIN)* (version 2.6.70)^42^. Interaction data were filtered to contain only genes included in the full single-copy data set. An undirected interaction network was visualized and analyzed using the *igraph* package (http://igraph.org) in *R*. Each node in the graph represents a single-copy nuclear gene family while each edge in the graph indicates a physical interaction in Arabidopsis. Nodes were colored by gene tree topology and diameter of nodes are proportional to bootstrap support values for the gene tree (see Fig. S2a-c).

We asked if genes displaying the same topology are clustered with each other in the network by calculating nominal assortativity^68^. Assortative mixing/clustering of gene tree topology results across the network was quantified by the assortativity coefficient (*A)* of the network. Positive *A* indicates clustering of genes with the same topology, while negative *A* indicates over-dispersal. We calculated the observed *A* for our network as well as a null distribution of *A* generated by randomly assigning a topology to nodes in 10,000 replicates of our network.

#### Mapping of gene coordinates to *A. thaliana* and *C. rubella* nuclear genomes

Topology results were mapped to the nuclear genome of *C. rubella* using the gene coordinates from the GFF file associated with the genome assembly. Genome maps were visualized using the *R* package, *Sushi*^69^, made available through *Bioconductor*^70^. Colored horizontal lines indicate genes displaying each topology. The length of each line represents the bootstrap support value found at the topology-defining branch in the gene tree (see Fig. S2a-c).

#### Detection of linkage disequilibrium

Topology results mapped to the *C. rubella* genome were used to ask if genes displaying the same topology are clustered together linearly along chromosomes. We assessed the physical clustering of A(BC), B(AC), and C(AB) genes with two measures: 1) number of instances in which genes with the same topology are located within 10kb of each other (Fig. S6a), and 2) number of instances in which neighboring genes share topology, regardless of distance (Fig. S6b). We established a null distribution for both measurements by generating 10,000 maps of the *C. rubella* genome in which observed location of single-copy genes and the overall gene tree frequencies were maintained, but the assignment of topologies to genes was randomized across chromosomes. Measure 1 and measure 2 were calculated for each of the 10,000 replicates to obtain null distributions.

#### Statistical Analyses

All statistical tests were performed in *R* (version 3.4). Below, we describe methods used to assess the significance of our results. Our general strategy was to provide sufficient information to enable readers to make their own interpretations of the data; toward that goal, we have included Bonferroni corrected and uncorrected (raw) *p-*values for each experiment where corrections could be applied (Tables S5 and Table S5 or within supplemental text). The conclusions we draw are statistically robust, and thus are not affected by whether significance is assessed by raw or Bonferroni corrected *p-*values. The fact that the majority of the *p-*values in support of our conclusions are significant shows that we are not ‘cherry picking’. Thus, our results are unlikely to have been affected by type-one error that can be associated with multiple tests. Therefore, in order to avoid inflation of type-two error, we report raw *p*-values in the main body of the manuscript.

#### *D, F,* and *D_GT_*-statistics

We calculated *D*, *F*, and *D_GT_* for both the full single-copy and conservatively single-copy data sets. Confidence intervals were obtained by resampling either dataset to generate 10,000 bootstrap replicates, recalculating *D/F/D_GT_* for each replicate. The resulting distributions were compared using the Z-test. To account for potential autocorrelation bias caused by non-independence of linked genes, *D/F/D_GT_* were also calculated using block bootstrapping. For *D* and *F*, block bootstrapping was achieved by simply bootstrap resampling from the available gene alignments and recalculating *D/F* with each replicate. For D_GT_ block bootstrapping was accomplished by splitting the dataset into 100 equal size blocks of neighboring genes based on position along *C. rubella* chromosomes. Blocks were then bootstrap resampled 10,000 times and *D_GT_* was recalculated with each replicate to obtain a distribution*. P*- values from analyses of the whole genome were Bonferroni adjusted for four comparisons for *D_GT_*.

#### Phylogenetic network reconstruction and introgression analysis

*PhyloNet* models were statistically compared by calculating AIC and BIC scores for each tree/network with the following expressions:

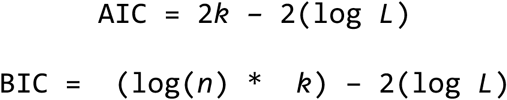

where *k* is the number of free parameters in the model, *n* is the number of input gene trees, and *L* is the maximum likelihood value of the model. We compared hypotheses by calculated difference in AIC and BIC scores for each given tree/network relative to the most likely network (ΔAIC and ΔBIC).

#### Node depth based test of species branching order

Frequency distributions of node depths were plotted. Two-tailed *T*-tests and Wilcoxon rank sum tests were performed to assess differences in distribution means and medians, respectively. *P*-values were Bonferroni corrected for six comparisons.

#### Divergence based test of IG directionality

Frequency distributions of node depths were plotted. Two-tailed Wilcoxon rank sum tests were performed to assess differences in distribution medians. *P*-values were Bonferroni corrected for three comparisons.

#### GO category enrichment

Enrichment of GO categories was assessed by comparing GO categories of A(BC) genes versus B(AC) genes. For each GO category, two-by-two contingency tables were constructed and used to perform Fisher’s exact tests. Results from two-tailed and one-tailed tests are reported. *P*-values from primary comparisons were Bonferroni corrected for three comparisons.

#### Protein-protein interaction network

Clustering in the interaction network was quantified with an assortativity coefficient (*A*)^68^. To assess significance of the observed *A*, we randomly assigned one of the three topologies (keeping the frequency of each topology the same as in the original data set) to genes in 10,000 copies of the network. We computed *A* for each of the 10,000 networks to obtain a null distribution of *A* and used the null distribution to perform a two-tailed *Z*-test.

#### Haplotype block linear clustering

We quantified linear clustering of topologies by counting the number of occurrences of proximal and neighboring genes in the observed data. We assessed the significance of the observed values by generating null distributions from 10,000 datasets in which the topologies were randomized. We used the null distributions to perform two-tailed *Z*- tests. *P*-values were Bonferroni corrected for six comparisons.

#### Data Availability

Gene tree data are linked to the online version of the paper. Scripts and input files used to perform analyses are available at: https://github.com/EvanForsythe/Brassicaceae_phylogenomics.

**Supplementary Information** is linked to the online version of the paper.

## Acknowledgments

The data reported in this paper are provided in the Supplementary Information. Scripts used to perform analyses are available at: https://github.com/EvanForsythe/Brassicaceae_phylogenomics. This work was funded by NSF grants 1409251, 1444490, and 1546825 to MAB. We thank M. J. Sanderson, M. M. McMahon, E. Lyons, D.B. Sloan, M.P Simmons, R. N. Gutenkunst, A. E. Baniaga, and S. M. Lambert for helpful discussions and M. T. Torabi, M. C. Borgstrom, and D. S. Clausen for statistical consultation. Finally, this work benefited greatly from input of the PaBeBaMo research group in the School of Plant Sciences, University of Arizona.

## Author Contributions

E.S.F and M.A.B conceived the study. A.D.L.N performed organellar genome assembly. E.S.F performed all other analyses. E.S.F and M.A.B wrote the manuscript with input from A.D.L.N. All authors approved of manuscript before submission.

## Author Information

The authors declare no competing financial interests. Correspondence and requests for materials should be addressed to M.A.B. at.

